# Positive germline selection of mtDNA: evidence from the oocyte

**DOI:** 10.64898/2025.12.31.697248

**Authors:** Melissa Franco, Zoë Fleischmann, Sofia Annis, Auden Cote-L’Heureux, Dylan Aidlen, Mark Khrapko, Benjamin Vyshedskiy, Mary K. Hartigan, Daniil Mirzoyan, Jacob Bandell, Konstantin Popadin, Dori C. Woods, Jonathan L. Tilly, Konstantin Khrapko

## Abstract

Purifying selection is the removal of detrimental mutations. Nuclear DNA mutations are purged by removing mutant individuals, embryos, or germline cells. Because mtDNA is present in numerous copies per cell, purifying selection of mtDNA mutations introduces an additional layer of complexity. The intracellular mutant fraction can change due to clonal expansion/reduction of mutant mtDNA molecules in germline cells. Different research groups have reported either negative (purifying), or positive (destructive) germline mtDNA selection. In this study, we use recently published high-fidelity data on mtDNA mutations in individual human oocytes from the Makova laboratory (Arbeithuber et al., 2025) to resolve this disagreement.

To assess selection, we used the “selective expansion” metric ***S***_***t***_, which compares the average weighted rate of clonal expansion in a set of mutations tested for selection (e.g. non-synonymous mutations) to that of the set of synonymous mutations. We found that, on average, noncoding oocyte mutations in the control region expand faster than synonymous mutations, i.e., are under positive selection. Intriguingly, mutations in the coding region, the detrimental ones in particular, were also on average under positive selection. The prescence of average positive selection does not preclude purifying selection against individual mutations; it only indicates that positive selection dominates in aggregate mutational dynamics. Positive selection impacts the dynamics of *de novo* germline mutations that arise in primordial germ cells or oocytes and have not yet been inherited by the next generation. Contrastly, mutations that are passed to subsequent generations are under average purifying selection. We suggest that initial positive selection begets subsequent purifying selection by increasing intracellular mutant fractions to levels at which their deleterious effects become phenotypically apparent and can be efficiently removed.

## Introduction

Mutational pressure on mtDNA exceeds that of nuclear DNA by over two orders of magnitude per nucleotide (Khrapko et al., 1997). Moreover, mtDNA mutations may be particularly prone to Muller’s Ratchet (Bergstrom and Pritchard, 1998), (Popadin et al., 2007). It’s therefore not surprising that mtDNA mutations must be kept under stringent purifying selection. Indeed, nascent mtDNA mutations that “bombard” mtDNA are mostly nonsynonymous (nonsynonymous/synonymous ratio is approximately 3:1). However, ancient mtDNA mutations that have existed in humans for thousands of generations (such as mutations in deep phylogenetic branches connecting modern and archaic humans), have a nonsynonymous/synonymous ratio of about 1:10, approximately 30x lower than the ratio in the nascent, “bombarding” mutations. Thus, selection should ultimately remove 29 of 30 nonsynonymous mutations.

Classical purifying selection typically proceeds through reduced survival and/or reproductive success of carriers of deleterious mutations. In contrast, *germline* purifying selection can act earlier by eliminating mutant germline cells rather than mutant individuals. Such early selection may reduce the developmental, reproductive, and energetic costs associated with producing severely affected offspring that would later be removed by organismal-level purifying selection.

Mitochondrial germline selection involves an additional layer of complexity. Because each cell contains many mtDNA molecules, selection on mtDNA mutations includes not only the removal of mutant germline cells, analogous to nuclear germline selection, but also changes in the proportions of mutant mtDNA molecules within individual cells. These intracellular shifts reflect the expansion or contraction of mutant mtDNA clones. Adding to complexity is that clone dynamics are driven not only by selection but also by random genetic drift within the intracellular mtDNA population (Coller et al., 2001).

Although purifying selection of mtDNA mutations is a well-established phenomenon, its mechanisms in the germline remain incompletely understood. A seminal study (Stewart et al., 2008) demonstrated purifying germline mtDNA selection in mtDNA mutator mice, as evidenced by elevated synonymity among inherited de novo mutations, a finding recently confirmed (Kremer et al., 2025). However, mutator-mouse-derived mutations used in these experiments differ substantially from natural mtDNA mutations in their spectrum and synonymy.

In humans, an influential study (Floros et al., 2018) initially proposed purifying germline selection in primordial germ cells (PGCs), based on an apparent increase in the synonymity of mtDNA mutations through PGC development. This result appeared to be supported by studies showing that, in humans, known mtDNA variants are more likely to be transmitted than unknown, i.e., more likely detrimental (Li et al., 2016), (Wei et al., 2019). However, we later showed that the apparent synonymity increase in PGCs resulted from co-amplification of a nuclear mtDNA pseudogene (NUMT). Importantly, the authors also published single-cell PGC mutational data that were not affected by NUMT contamination (Floros et al., 2022). Our analysis of these single-PGC data demonstrated preferential expansion of nonsynonymous mutations, providing evidence for positive germline selection in PGCs (Fleischmann et al., 2021), (Fleischmann et al., 2024).

Very recently, a high-fidelity single-cell mtDNA mutational analysis of oocytes was published (Arbeithuber et al., 2025). In apparent agreement with Floros et al., they reported purifying germline selection, especially at higher mutant fractions. This conclusion challenges our inference that germline selection in PGCs was positive: some ‘carryover’ of that positive selection is expected in oocytes that are derived from PGCs. To resolve the inconsistency, we reanalyzed the Arbeithuber et al. data using a specially developed selective expansion analysis approach. We found that the impression of purifying selection on coding mutations in oocytes arose from comparison with non-coding mutations, which are themselves highly positively selected. After correction, the prevailing selection pattern among nascent coding mutations turned out positive.

Importantly, our finding of positive selection in PGCs and oocytes does not necessarily contradict reports of purifying germline selection by Stewart et al. In the latter study, selection is estimated for mutations that have passed several generations, whereas PGC mutations in Floros et al. and in our study are nascent germline mutations that have not yet been transmitted to the next generation. Thus, we expect that early, predominantly positive, germline selection is ultimately surpassed by negative selection, which operates on a longer timescale.

## Results

### S_t_: a selection statistic based on threshold-crossing expansion

To quantify selection, we relied on direct analysis of clonal expansion of mutants rather than on the conventionally used indirect dN/dS synonymity metrics. dN/dS-style approaches are biased by the strong interdependence between synonymity and the mutational spectrum in mtDNA (manuscript in preparation). Clonal expansion is central to mtDNA inheritance and evolution. To be fixed in the maternal lineage and the species, the mutation must gradually expand from a single nascent mutant molecule to homoplasmy (100%) in the germ cell. Intercellular expansion (or decline for negative/purifying selection) is driven both by random genetic drift and selection within the cell’s mitochondrial populations. At the intercellular level, clonal expansion/decline is further influenced by differences in mutant cell proliferation and by their attrition. Intercellular dynamics is also driven by the superposition of drift and selection in the cellular population. Thus, to measure germline selection, we need to separate it from random drift.

By definition, molecular selection implies that the expansion of mutant DNA molecules *reproducibly* exceeds (or lags behind) that of neutral variants. Reproducibility is critical because mutations commonly outpace one another due to random drift; only by showing reproducibility can selection be demonstrated. However, in relatively small single-cell mtDNA mutational datasets, including (Arbeithuber et al., 2025) – (70 oocytes), most *de novo* mutations yield a single observation each, precluding demonstration of reproducibility for individual mutations. In this case, selection must be inferred from the average expansion across multiple mutations. Thus, we choose a set of mutations that we wish to test for selection, which can be, e.g., coding, non-coding, or non-synonymous mutations (“set A”). Selection is established if expansion in set A, on average, is significantly faster (or slower) than among reference mutations assumed not to be under selection (e.g., synonymous mutations). Importantly, reference mutations must originate from the same cells and the same experiment as the tested mutations. If so, the dynamics of random drift are identical in the tested and reference sets, so any differences between them can be attributed to selection in the tested set. Then, subtracting the average expansion in the synonymous set from that of the set A gives an estimate of selection in the set A. Below, we formalize this intuitive approach.

To quantify the progression of mutations of a given set A, starting from single nascent mutant molecules to expanded mutant clones, we first define ‘expanded mutations’ as mutant clones with an intracellular mutant fraction above threshold ***t***. Then, a ‘natural’ measure of expansion would be ***E***_**t**_**(*A*)**, i.e., the share of aggregate mutant fraction of expanded mutations (MF> ***t***) among all mutations of the set A. The schematic below illustrates this concept, showing an imaginary example of positive selection. The exaggerated high-fraction shoulder of set A illustrates how ***E***_**t**_ is effective in measuring differences in expansion between sets of mutations.

To quantify the selection-only portion of expansion, synonymous expansion (driven by random drift) is subtracted from the expansion of mutations of set A:

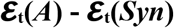

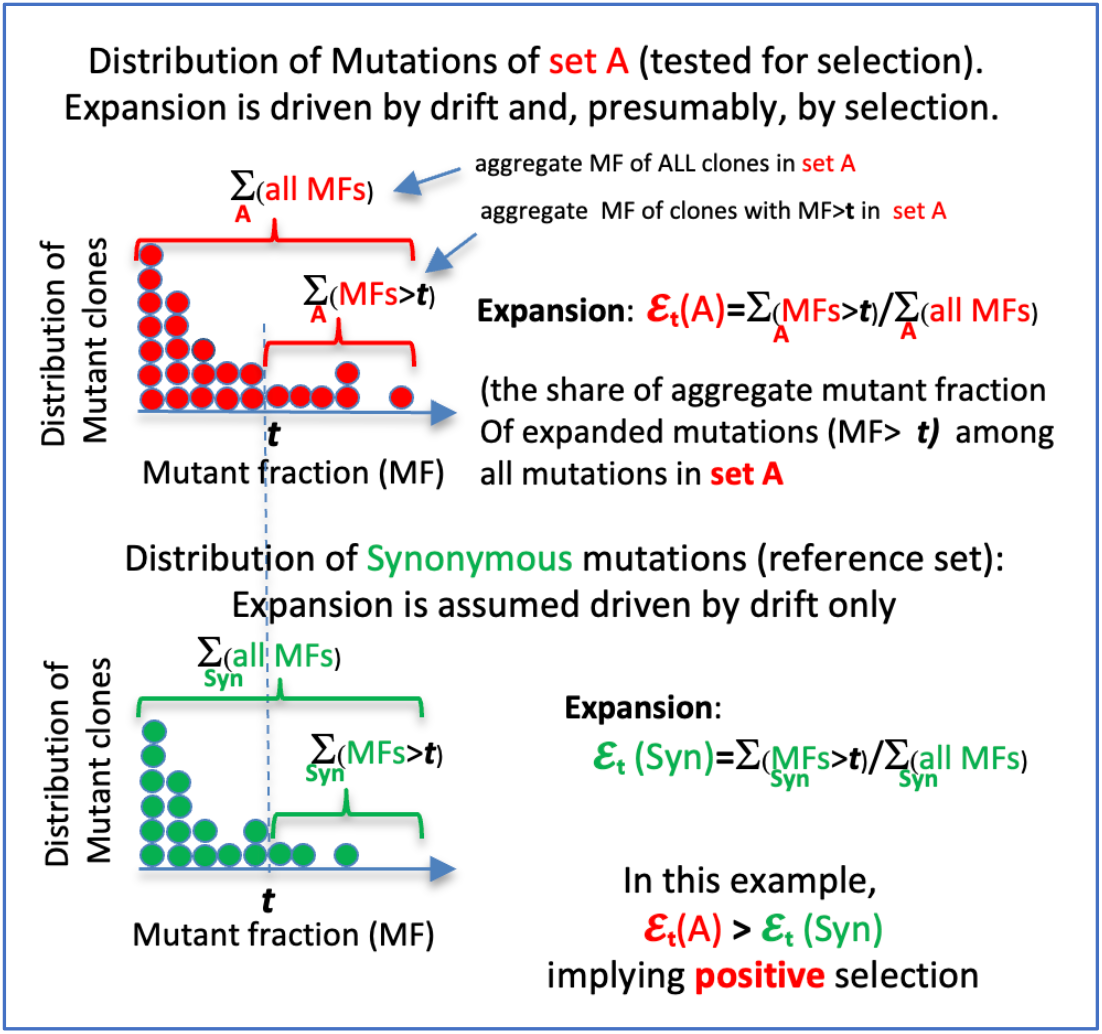

This quantity represents the portion of expansion attributable to selection. To obtain a universal normalized scalar measure of selection, this “selective expansion” is divided by the neutral expansion of synonymous mutations, defining the selection statistic ***S***_t_:

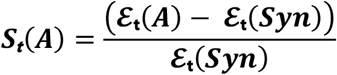

***S***_t_ represents the contribution of selection-driven expansion relative to expansion driven by random drift. Thus, ***S***_t_ > 0 indicates positive selection, whereas ***S***_t_ < 0 indicates negative/purifying selection. ***S***_t_ ~ 0 implies neutral dynamics similar to those of synonymous mutations. ***S***_t_ ~ 1 indicates that the contribution of selection is comparable to that of random drift.

Statistical significance of selection was evaluated by Monte Carlo resampling of subsets from the set of synonymous mutations and calculation of the fraction of simulations in which simulated ***S***_t_ values were equal to or more extreme than the experimentally observed ***S***_t_, yielding a one-sided p-value. We then repeated ***S***_t_ analysis for a range of thresholds ***t*** to confirm the stability of the result (see **Methods** for details).

### Non-coding mtDNA mutations are under strong positive selection in oocytes

We first asked whether non-coding oocyte mutations were under positive selection. If non-coding mutations are indeed under positive selection, this may create an artificial impression of purifying selection in coding mutations, since they were compared to non-coding mutations as a reference by Arbeithuber et al. (**Supplementary Note**). We and others (Nekhaeva et al., 2002), (Korotkevich et al., 2025) have previously observed positive selection of non-coding (d-loop) mutations in single somatic cells; thus, this possibility was biologically plausible. We started by calculating ***S***_***t***_ at ***t***=0.01, following Arbeithuber et al.’s choice of the threshold. The results (**Table 1**) indeed demonstrate highly significant (p~0.002) and strong (***S***_***0.01***_ = 0.36) positive selection for non-coding mutations.

**Table 1.**

| Mutation Type | $\mathcal{S}_{0.01}$ | $\mathcal{S}_{0.01}$ P-value |
| --- | --- | --- |
| Non-coding / d-loop | 0.36 | 0.0024 |
| Coding w/o syn | 0.14 | 0.13 |
| Coding conservative | 0.25 | 0.028 |

To demonstrate the robustness of this result, we additionally scanned thresholds of 0.003 and 0.001 and showed that both indicate positive selection of noncoding mutations (**Table S1**). Note that a decrease of ***S***_***t***_ at lower t is unsurprising: as ***t*** decreases, the tested set increasingly includes weakly selected and neutral mutations, reducing the average signal of selection.

### Coding oocyte mutations are under “destructive” positive selection

Strong positive selection of non-coding mutations implies that the apparent purifying selection of coding mutations reported by Arbeithuber et al. arises from using a positively selected reference (non-coding mutations), which biases comparisons toward a superficial deficit of expanded coding mutations. To assess and correct the bias, we compared the average expansion of coding mutations (including RNA mutations but excluding synonymous mutations) to that of synonymous mutations, i.e., calculated the corresponding ***S***_***t***_ (**Table 1**). Intriguingly, this revealed not only the absence of purifying selection but also weak (***S***_***0.01***_ ~0.11) and nonsignificant (p ~0.15) *positive* selection.

Previous research in single somatic cells indicated that positive selection on mtDNA mutations can be destructive, i.e., favoring detrimental mutations (Nekhaeva et al., 2002), (Korotkevich et al., 2025). We therefore hypothesized that the trend towards positive selection observed in coding mutations exemplifies such destructive selection. If so, then focusing on the most detrimental mutations should strengthen the positive selection trend and potentially render it statistically significant.

To test this hypothesis, we used GERP++, an evolutionary conservation score (Davydov et al., 2010) (see **Methods**) as a proxy for the deleteriousness of mutations. More generally, such analysis helps address an essential conceptual question: is the positive selection of coding mutations driven, at least in part, by their detrimental effects? If more conserved coding mutations show stronger positive selection than less conserved ones, then positive selection is linked to deleterious mutations. Conversely, if the average selection among conserved mutations is not higher than in all mutations, then the hypothesis is likely incorrect.

To discern these possibilities, we selected the top ~2/3 most conserved coding mutations (i.e., GERP>10) and repeated ***S***_***0.01***_ analysis. As shown in **Table 1**, for conserved coding mutations, the intensity of selection increased and the result became statistically significant. This effect remains robust across all three thresholds tested (**Table S1**). This result supports the (perhaps counterintuitive) idea that positive selection of coding mutations in the germline is linked to their detrimental effects and thus, in this sense, is “destructive”.

### Visualization of selection with cumulative curves

To visualize and compare mutation expansion in different sets of mutations across the full range of thresholds ***t***, we constructed cumulative expansion curves (**Figure 1**). These curves are conceptually similar to those previously used to analyze cancer mtDNA mutations (Yuan et al., 2020) and provide an approximate graphical representation of selection throughout the range of mutant fractions. To construct a curve, for each mutation in a set, we calculated the aggregate mutant fraction as the fraction of mutations with MFs higher than the MF of that given cell, divided by the aggregate MF of all mutations. In other words, we calculated the value of ***E***_**t**_ using this mutation’s MF as the threshold ***t. E***_**t**_ values for all mutations of the set were plotted as a function of ***t***.

**Figure 1.**
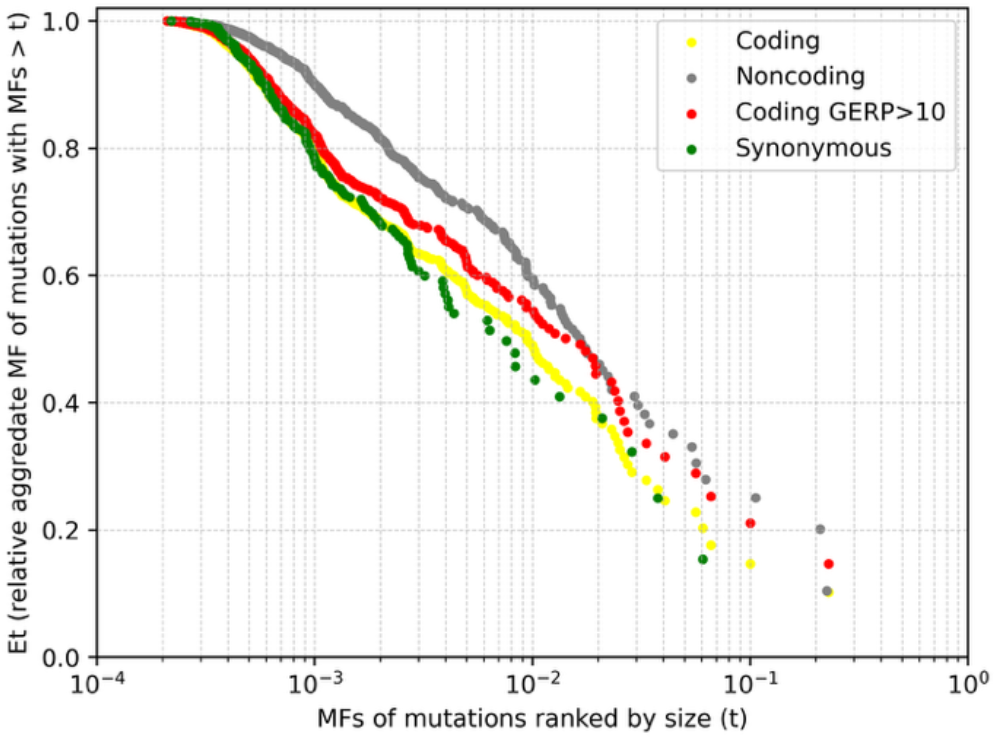
An overview of the dynamics of the expansion of the different types of mutations that offers a visual representation of selection. A relative shift of a curve upwards from the synonymous green curve represents an excess of expansion over random drift at a given ***t*.**Shift up implies positive selection: it means that a larger proportion of mutations exceed the threshold ***t*** than expected from random drift. Dashed lines indicate points where ***S***_***t***_ and Monte-Carlo p-values (Table 1 and S1) were calculated.

In this representation, upward displacement of a curve relative to the synonymous curve indicates an excess expansion beyond neutral drift expectations (positive selection). Comparison of the curves qualitatively illustrates selection from a broader perspective than the quantitative 3 individual ***S***_***t***_ calculations in Tables 1 and S1, whose positions are indicated by vertical dashed lines. The curves are merely a convenient tool to qualitatively represent the consistency of the selection patterns and the general distributions of the data. The particular local features of these curves (such as points where they diverge, drop-offs, and gaps are random perturbations of the data and should not be considered important.

### Average positive germline selection reflects a balance between positive and negative selection

The average measures of selection used in this study are suited for identifying average selection patterns in sets of mutations (e.g., all non-synonymous or coding mutations), but cannot characterize the effects acting on specific mutations. Thus, individual mutations within a positively selected set may themselves be neutral or even negatively (purifyingly) selected, but their contribution may be masked by the overwhelming signal from positively selected mutations. This tentatively suggests that some germline mutations, including those not yet inherited, are under purifying selection, but these mutations do not dominate the overall selective landscape.

### Tests for potentially missed negative selection: dominant-lethal mutations and comparison with the numerical approach

Measuring negative selection is, as expected, more challenging than measuring positive selection. Negatively selected mutations are expected to remain at low fractions or disappear altogether. Hypothetically, under extreme purifying selection modes, a particularly aggressive mutation (e.g., causing a lethal burst of ROS) could trigger a rapid elimination of the mitochondrion or cell in which it arose and therefore of mutation itself (which we denote as a “dominant lethal” mutation). Such dominant-lethal mutations cannot be measured directly. Nevertheless, dominant lethal selection could be inferred by searching for “holes” in mutational spectra, analogous to the classical survival-bias analysis of the returning WWII warplanes. Our preliminary observations suggest that such “holes” may indeed exist in the oocyte mutational spectrum, but establishing the significance of this finding requires more work and larger datasets. This type of negative selection cannot be readily detected by our “selective expansion” analysis or any analysis based on direct measurement of mutational dynamics. Although this selection could potentially be detected by indirect approaches, e.g., dN/dS, such approaches are biased due to the strong interdependence of synonymity and mutational spectrum in mitochondria (manuscript in preparation).

Another potential problem is the difference between the approaches used in our analyses and in previous studies. We used a weighted approach that correctly accounts for the mutant fraction. In contrast, Arbeithuber et al. used a conventional numerical approach that underweights highly expanded mutant clones and may overemphasize the selection of smaller mutations (see **Methods** for details). Thus, to account for the difference in the approaches, we also used the numerical version of ***S***_***t***_, denoted ***S***^**n**^_***t***_ (Supplementary **Table S2)**. A comparison of **Table S1** and **Table S2** shows that the numerical approach detects no negative or purifying selection, confirming our conclusion that there is no purifying selection. As far as positive selection is concerned, it is substantially reduced and no longer statistically significant when quantified using the numerical approach. This implies that destructive positive selection is primarily dependent on the increased intracellular expansion of mutant clones rather than on a larger number of clones (additional details can be found in the captions for **Table S1** and **Table S2**).

## Discussion

### Destructive positive germline selection: new, high-fidelity evidence from oocytes

This study sought to reconcile our previous finding of positive selection in PGCs, the precursors of oocytes (Fleischmann et al., 2024), and the purifying selection signatures of oocyte mutations reported by Arbeithuber et al., 2025. Arbeithuber et al. used a technically superior methodology of mutational analysis, double-stranded sequencing (Kennedy et al., 2013), which they adapted for single-cell analysis (Arbeithuber et al., 2020). This approach drastically reduces PCR-related errors and therefore yields high-quality mutational data. These data provided an important opportunity to test our conclusions about positive germline selection based on PGC mutational data.

We noted that Arbeithuber et al. primarily assessed purifying selection based on the relatively lower prevalence of coding mtDNA mutations in comparison to non-coding mutations. We therefore asked whether positive selection of non-coding mutations, rather than negative/purifying selection of coding mutations, could explain their observations. Our interpretation was supported by previous demonstrations that non-coding mutations are indeed prone to positive selection in single cells, at least *in somatic tissues* (Nekhaeva et al., 2002), (Korotkevich et al., 2025). With this hypothesis in mind, we devised a metric of selection, ***S***_***t***_, that uses neutral synonymous mutations, rather than non-coding mutations, as a reference. Using this approach, we observed strong positive selection for non-coding mtDNA mutations in the Arbeithuber et al. dataset (**Tables 1** and **S1**).

If non-coding mutations are under positive selection, they cannot be used as an unbiased reference for assessing selection of coding (or any other) mutations. We therefore reanalyzed the potential selection of coding mutations using the neutral metric, ***S***_***t***_. As expected, ***S***_***t***_ provided no evidence for purifying selection of coding mutations. Moreover, ***S***_***t***_>0 suggested that, as a set, coding mutations may expand faster than synonymous mutations, i.e., they had a trend toward *positive* selection, though this observation was not statistically significant (**Table 1)**.

Importantly, we additionally found that the effect of positive selection is stronger and becomes statistically significant among the most conserved (i.e., most likely detrimental) coding mutations. In other terms, mtDNA selection in the germline is, at least in part, “destructive”. Finally, our observation of overall positive selection does not imply the absence of purifying selection. There are clear examples of pathogenic mtDNA mutations undergoing purifying selection in the germline (Fan et al., 2008), (Freyer et al., 2012). Rather, our results most likely mean that the average positive-selection signal observed in oocyte mutations reflects a balance between stronger or more prevalent positive selection acting on some mutations and weaker or less prevalent purifying selection acting on others.

### “Destructive” and “selfish” positive selection in mtDNA: what is known

While positive selection in mtDNA, and especially “destructive” selection of detrimental mutations, seems conceptually counterintuitive, it is consistent with several prior reports, both in somatic cells and in the germline. In somatic cells/tissues, positive selection has been observed in cardiomyocytes (Nekhaeva et al., 2002), (Korotkevich et al., 2025), skeletal muscle (Bua et al., 2006), (Nicholas et al., 2009), *substantia nigra* (Kraytsberg et al., 2006), cancer (Yuan et al., 2020), and fibroblasts (Korotkevich et al., 2025), to mention just a few studies. Positive destructive selection in whole somatic tissues has been reported too (Li et al., 2015), (Kuiper et al., 2025). Most importantly, positive selection of detrimental mutations has been shown to act in the germline during transmission to the offspring as well (Otten et al., 2018), (Franco et al., 2020), (Zhang et al., 2021), (Franco et al., 2023). The possible mechanisms of positive selection of detrimental mtDNA mutations have been widely discussed since the seminal paper by Aubrey de Grey (de Grey, 1997), who proposed the SOS (“survival of the slowest”) model of positive selection. Multiple cases and mechanisms of positive selection of detrimental mutations have been proposed since then (e.g., see SSD – Stochastic Survival of the Densest (Insalata et al., 2022) for a recent one).

Positive selection of *noncoding* mutations is often viewed as a selfish mode of selection, in which mutations expand by upregulating their own replication, presumably through alteration of regulatory sequences. Selection of noncoding mutations has been proposed to be driven by either the replication/transcription switch associated with G-quadruplex structural polymorphism ((Wanrooij et al., 2010), (Tan et al., 2016), (Agaronyan et al., 2015), (Gupta et al., 2023) or by the second light strand promoter (Tan et al., 2022). These two mechanisms may correspond to two blocks of mutations associated with intracellular mtDNA expansion identified by Nekhaeva et al. (2002), the C-tract block and the 16028-16050 block (Nekhaeva et al., 2002).

### Selection of not-yet-inherited vs. inherited mutations

There is no doubt that a major force of mtDNA evolution is purifying selection, which appears at odds with our finding of positive selection in the germline. There is no contradiction, though. Note that studies showing purifying selection in mutator mouse mtDNA, i.e., (Stewart et al., 2008), (Kremer et al., 2025), in human mother-child pairs (Li et al., 2016), (Wei et al., 2019), 4-generation groups (Liu et al., 2022), and over a wide evolutionary scale (Cavadas et al., 2015), all are dealing with *inherited* mutations that have passed one or more generations, whereas positive selection reported in this study pertains to *de novo, not-yet-inherited* ones. This implies an interesting possibility: the prevalent mode of selection may change for those *de novo* germline mutations that survive and cross the intergenerational boundary and become ‘inherited’.

Not-yet-inherited mutations analyzed here are measured in oocytes, but they could have been generated throughout germ-cell development, including PGCs. We have previously demonstrated positive selection of mtDNA mutations in single PGCs (Fleischmann et al., 2024). Because oocytes are direct descendants of PGCs, the positive selection pattern we now observe in oocytes should involve selection in PGCs and any subsequent selection occurring while PGCs develop into mature oocytes, including the enigmatic syncytial cyst stage (Lei and Spradling, 2016). The PGCs, the syncytium, and non-dividing developing oocytes represent distinct cellular environments, and the mechanisms and the dynamics of selection may differ across these stages. Our preliminary observations comparing mtDNA mutations in PGCs vs. oocytes suggest an episode of purifying selection removing the most extensively expanded nonsynonymous PGC mutations (**manuscript in preparation**) and likely partially erasing the signature of positive PGC selection of oocyte mutations. It is thus possible that the positive selection we observe in oocyte mutations includes a residual signature of PGC selection. Some positive selection may also occur in developing oocytes: positive selection of a specific detrimental mutation during oocyte maturation has been reported in the mouse (Zhang et al., 2021).

### The potential role of the “destructive” germline selection

Positive germline selection of detrimental mutations appears counterproductive, because germ cells with expanded detrimental mtDNA mutations (or their mutant descendants, from embryos to adults) are likely to eventually terminate as a lineage. We are confident about this because, as argued in the Introduction, an absolute majority of non-synonymous bombarding mutations (~29 of 30 that are not removed by the drift) are not fixed in the population, and therefore must be selectively eliminated one way or another. We hypothesize that positive selection may present a mechanism that allows carriers of such mutations to be eliminated earlier than later, as a higher fraction of a detrimental mutation usually makes the defect more pronounced. Then mutation carriers can be promptly removed, thereby sparing the mother the risk of carrying and raising offspring that will fail at a later developmental stage, in adulthood, or will fail to produce viable future generations (this may happen generations later further increasing cumulative burden). This may not necessarily be an adaptive mechanism, but at least this may be a reason why nature did not invest in eliminating destructive selection.

Of note, the positive selection/elimination scenario would be useless if positive selection started to operate at very low mutation fractions, such as those of recently generated mutations. This would be equivalent to merely increasing the mutation rate. In contrast, if positive selection were initiated at a higher mutant fraction, it would target specifically the carriers of the break-through, expanded mutant clones, which were “lucky enough” to benefit from random drift which pushed them over the selection threshold, where they were expanded further by selection. Interestingly, positive germline selection indeed appears to be initiated at higher mutant fractions: our study of mother–child inheritance of detrimental mutations reveals an arch-shaped selection profile, in which selection is absent or even negative at low mutant fractions, but becomes positive at intermediate fractions, and then reverts to negative at high (likely unsustainable) levels (Franco et al., 2023). This wave-shaped selection profile appears to be ideally positioned to enhance purifying selection of expanding mutations.

In conclusion, we have demonstrated the overall positive germline selection among first-generation, not-yet-inherited germline mutations. This, however, does not mean the absence of negative selection. Our results support a model in which mtDNA selection reflects a dynamic interplay between early mostly positive (“destructive”) selection, which drives the expansion of recently generated mutations, and subsequent predominantly purifying selection. Destructive and purifying selection appear to jointly shape the fate of mtDNA mutations.

## Methods

### Data

Mutational data were retrieved from the Supplemental material variants.xlsx (Arbeithuber et al., 2025).

### Not-yet-inherited vs. inherited germline mutations

The mutations discussed in this study are those created in the current generation (unless otherwise indicated). We define current-generation mutations as those observed in no more than one oocyte from a given donor. Being non-repeated among oocytes from the same donor implies that the mutation was *unlikely* to be present at a significant fraction in the mother-oocyte that eventually developed into that oocyte donor; otherwise, it would likely appear in more than one of her oocytes. Thus, these non-repeated mutations were likely generated *after* the mother-oocyte developed into an embryo, implying that they were not inherited from the previous generation. On the contrary, mutations that have been repeated at least in two oocytes or an oocyte and a somatic tissue of the same individual were considered inherited from the previous generation because it’s unlikely that a nascent mutation gains enough prevalence early enough to be shared in this way. This definition of inherited mutations differs slightly from that of Arbeithuber et al., it is a less inclusive definition of the not-yet-inherited mutations.

### Weighted vs. numerical: definitions and justification

Mutant fraction (MF) is defined in this study as the fraction of mutant mtDNA molecules among all mtDNA molecules in a given pool of molecules. This pool can consist of molecules in a single cell, molecules in a tissue sample, or any unbiasedly pooled molecular data. This universal definition applies equally to an individual mutation or to a set of mutations (e.g., a set of coding mutations). This measure is the *weighted* mutant fraction because each mutation’s contribution to the aggregate MF is weighted proportionally to its individual MF. Note that each not-yet-inherited germline mutation in a single-cell mutational dataset represents an intracellular *clone* of mutant molecules. Thus, in the weighted mutant fraction approach, each mutation’s contribution to the aggregate mutation load is proportional to the clone size.

An alternative approach, which we dubbed the *numerical* approach, counts each mutation (i.e., particular nucleotide changes in a particular genomic position) once per cell, implying that the initial mutational event that caused that mutation happened once in this cell or the upstream cellular lineage (and survived genetic drift). The numerical approach is conventionally used in mutational analysis, and was used by Arbeithuber et al. In our view, the numerical approach is the method of choice in a selection study. Numerical MF is insensitive to the size of intracellular mutant clones (large and small clones make the same contribution), and therefore to the intensity of the intracellular selection, which is responsible for the systematic part of the differences between large and small clones.

Furthermore, the weighted distribution of germline mtDNA mutations better approximates the probability of inheritance than the numerical distribution. Indeed, the probability of being inherited is not a fixed quantity per mutation present in the oocyte but rather is proportional to the *fraction* of mutant molecules because the population of mtDNA in the cell is under random genetic drift. The probability of any subpopulation to survive the drift (and therefore get inherited) in proportional to the fraction of the subpopulation. If a mutation is under selection, this adds an additional multiplicative factor to the probability of inheritance (>1 for positive and <1 for negative selection), but the basic proportionality to mutant fraction remains. In other words, the weighted approach is correctly focused on what’s most important for the germline mutations - their probability of inheritance.

To quantify selection, we use selective expansion ***S***_***t***_ which was described in the **Results** section. ***S***_***t***_ is based on the comparison of the loads of tested and synonymous mutations above and below a threshold ***t. S***_***t***_ can be calculated both in the weighted and numerical paradigm, depending on whether the mutant load is calculated as weighted or as numerical. Based on the justification above, we used the weighted approach in the mainstream of this study. However, for comparison, we also tested the numerical approach to calculate ‘numerical ***S***_***t***_’, denoted ***S***^**n**^_***t***_. The results for ***S***^**n**^_***t***_ *are* shown in **Table S2** and discussed in the **Supplementary Note**.

We note that numerical approach may, too, be somewhat responsive to intracellular selection. For example, if there is positive selection of coding mutations, then because a relatively larger *number* of (faster expanding) coding than synonymous clones cross the threshold, will be positive ***S***^**n**^_***t***_. However, ***S***^**n**^_***t***_ is less sensitive than regular weighted ***S***_***t***_ because it does not capture the faster expansion of coding clones relative to synonymous clones *after* they cross the threshold.

### Monte Carlo significance of the relative expansion selection

To test the statistical significance of observations of selection, a Monte Carlo simulation was used. The null hypothesis was that there was no positive selection in the tested set of mutations. To test the null, the set of synonymous mutations was randomly sampled with replacement to generate simulated tested sets of the same size as the experimental tested set and ***S***_***t***_ was computed for each simulation (50,000 in this study). The fraction of simulations in which ***S***_***t***_ was equal to or larger than that observed in the experimental test set was used as the Monte Carlo p-value.

### GERP: a mutation conservation score

We used GERP++ (Davydov et al., 2010) as a metric of mutation conservation and as a substitute for the PhastCons and MLC scores used by Arbeithuber et al. We prefer GERP due to our strong understanding of, and extensive experience with this score. We have previously constructed and validated a particular embodiment of the GERP++ score (Gunbin et al., 2017), which produces a range of scores between ~-80 for most non-conserved sites to ~+40 for most conserved sites in mtDNA. This continuous metric shows more detail in the mutation conservation landscape than the binary PhastCons score. GERP is similar to the mitochondrial local constraint (MLC). GERP is expected to be generally consistent with the MLC score. Indeed, we, like Arbeithuber et al., observed a general decrease in average GERP score above MF 0.01 (from 1.9 below to −3.9 above).

## Acknowledgments

The authors are grateful to Kateryna Makova and Barbara Arbeithuber (Penn State University) for helpful discussions, criticisms, and suggestions.

## Supplement

### Note: Using positively selected non-coding mutations as a reference creates an appearance of purifying selection

To estimate selection of coding mutations, Arbeithuber et al. compared the coding/noncoding ratio at MF >0.01 to that at MF <0.01. This approach is conceptually similar to our ***S***_*t*_ measure. However, for comparison, it uses non-coding mutations, which are themselves positively selected. Such a reference is strongly biased and shifts selection estimates toward negative values. Additionally, the use of numerical (unweighted) counts disregards any expansion of clones that happens above the 0.01 threshold. This expansion is integral to the selection process, and its omission further unfairly increases the perceived negativity of selection (compare **Table S1** and **Table S2**).

Of note, positive selection of coding mutations may appear to contradict Arbeithuber et al.’s observation that the conservation score MLC is lower in *all* oocyte mutations with MF > 0.01 compared to those with MF < 0.01, as if there were purifying selection. However, all mutations include coding and non-coding mutations. Non-coding mutations are far less conserved (average GERP −6.0) than coding mutations (average GERP +5.6). Therefore, the observed decrease in conservation at high MFs may be due not to purifying selection on coding mutations, but rather to an increase of the proportion of noncoding mutations due positive selection on the latter. To test this possibility, we examined the conservancy of coding mutations alone. As expected, the trend not only disappeared but even reversed: the average GERP score of coding-only mutations showed an increase from 5.5 at MF<0.01 to 13.3 at MF>0.01, in agreement with positive selection.

**Table S1.** is an extension of Table 1 (with added thresholds of 0.003 and 0.001), demonstrating the robustness of the conclusions, i.e., confirming that the low p-values in **Table 1** are not due to threshold selection. At ***t***=0.01, the evaluation focuses on selection among high-fraction mutations. Interestingly, ***S***_***t***_ typically decreases at low ***t***, suggesting that positive selection is stronger/more prevalent at higher MFs. Alternatively, or in addition, this effect may also be caused by the expected “crowding” of mutations prone to positive selection at high MFs. Untangling these factors is a focus of our current work.

| Type of Mutation | $\mathcal{S}_{0.01}$ | $\mathcal{S}_{0.01}$ P-value | $\mathcal{S}_{0.003}$ | $\mathcal{S}_{0.003}$ P-value | $\mathcal{S}_{0.001}$ | $\mathcal{S}_{0.001}$ P-value |
| --- | --- | --- | --- | --- | --- | --- |
| Non-coding/d-loop | 0.36 | 0.0024 | 0.26 | <1e-04 | 0.16 | <1e-04 |
| Coding w/o Syn | 0.14 | 0.13 | 0.06 | 0.1645 | 0.02 | 0.248 |
| Coding conserved | 0.25 | 0.0281 | 0.13 | 0.0246 | 0.06 | 0.0313 |

**Table S2.** shows the numerical version of the metric ***S***_*t*_, i.e., ***S***^n^_*t*_. The definition of ***S***^n^_*t*_ is different from that of ***S***_*t*_ in that aggregate mutant fractions are replaced by mutation counts. This change reduces the emphasis on intracellular expansion/contraction of clones. We performed the ***S***^n^_*t*_ analysis to ensure we had not missed any selection to which ***S***^n^_*t*_ could be more sensitive than ***S***_*t*_ is. In comparison to ***S***_*t*_ values in Table S1, the ***S***^n^_*t*_ values for coding mutations are closer to zero (i.e., demonstrating weaker selection), and the statistical significance was completely lost. This reconfirms that intracellular expansion of clones is, overall, a strong component of selection.

| Type of Mutation | $\mathcal{S}^n_{0.01}$ | $\mathcal{S}^n_{0.01}$ P-value | $\mathcal{S}^n_{0.003}$ | $\mathcal{S}^n_{0.003}$ P-value | $\mathcal{S}^n_{0.001}$ | $\mathcal{S}^n_{0.001}$ P-value |
| --- | --- | --- | --- | --- | --- | --- |
| Non-coding | 1.01 | <1e-4 | 0.74 | <1e-04 | 0.59 | <1e-04 |
| Coding w/o Syn | 0.03 | 0.4409 | -0.06 | 0.7636 | -0.01 | 0.6163 |
| Coding conserved | 0.16 | 0.2614 | 0.03 | 0.4249 | 0.05 | 0.2035 |

By contrast, for *non-coding* mutations, ***S***^n^_*t*_ are more positive and statistically more robust than ***S***_*t*_. This observation may indicate a different mechanism of selection for non-coding mutations. Their selection appears to depend more on the increased number of mutant clones than on their intracellular expansion. Taken at face value, this may seem to indicate that at least some of these mutations stimulate the proliferation of germline cells, so clones are multiplied as cells that carry them proliferate. This is unlikely, however, especially because Figure 1 suggests that positive selection of non-coding mutations begins at very low mutant fractions, which are not expected to have functional effects on the entire cell. It is more likely that molecule-autonomous positive selection (expected for control-region mutations) may be improving the survival of nascent mutant molecules that would otherwise have been removed by random drift. Such improved survival is equivalent to a wider bottleneck and would appear to an observer as a higher number of clones than for synonymous mutations, thereby specifically boosting ***S***^n^_*t*_. This process adds large numbers of small clones, which increases ***S***^n^_*t*_ more than ***S***_*t*_. This aspect of mutational dynamics remains hypothetical and warrants further exploration.

## References

Agaronyan, K., Morozov, Y.I., Anikin, M., Temiakov, D., 2015. Replication-transcription switch in human mitochondria. Science 347, 548–551. 10.1126/science.aaa0986

Arbeithuber, B., Anthony, K., Higgins, B., Oppelt, P., Shebl, O., Tiemann-Boege, I., Chiaromonte, F., Ebner, T., Makova, K.D., 2025. Allele frequency selection and no age-related increase in human oocyte mitochondrial mutations. Sci. Adv. 11, eadw4954. 10.1126/sciadv.adw4954

Arbeithuber, B., Hester, J., Cremona, M.A., Stoler, N., Zaidi, A., Higgins, B., Anthony, K., Chiaromonte, F., Diaz, F.J., Makova, K.D., 2020. Age-related accumulation of de novo mitochondrial mutations in mammalian oocytes and somatic tissues. PLOS Biology 18, 1–37. 10.1371/journal.pbio.3000745

Bergstrom, C.T., Pritchard, J., 1998. Germline bottlenecks and the evolutionary maintenance of mitochondrial genomes. Genetics 149, 2135–46.

Bua, E., Johnson, J., Herbst, A., Delong, B., McKenzie, D., Salamat, S., Aiken, J.M., 2006. Mitochondrial DNA-deletion mutations accumulate intracellularly to detrimental levels in aged human skeletal muscle fibers. Am. J. Hum. Genet. 79, 469–480.

Cavadas, B., Soares, P., Camacho, R., Brandão, A., Costa, M.D., Fernandes, V., Pereira, J.B., Rito, T., Samuels, D.C., Pereira, L., 2015. Fine Time Scaling of Purifying Selection on Human Nonsynonymous mtDNA Mutations Based on the Worldwide Population Tree and Mother-Child Pairs. Human Mutation 36, 1100–1111. 10.1002/humu.22849

Coller, H.A., Khrapko, K., Bodyak, N.D., Nekhaeva, E., Herrero-Jimenez, P., Thilly, W.G., 2001. High frequency of homoplasmic mitochondrial DNA mutations in human tumors can be explained without selection. Nature Genetics 28, 147–150.

Davydov, E.V., Goode, D.L., Sirota, M., Cooper, G.M., Sidow, A., Batzoglou, S., 2010. Identifying a high fraction of the human genome to be under selective constraint using GERP++. PLoS Comput Biol 6, e1001025. 10.1371/journal.pcbi.1001025

de Grey, A., 1997. A proposed refinement of the mitochondrial free radical theory of aging. Bioessays 19, 161–166.

Fan, W., Waymire, K.G., Narula, N., Li, P., Rocher, C., Coskun, P.E., Vannan, M.A., Narula, J., MacGregor, G.R., Wallace, D.C., 2008. A mouse model of mitochondrial disease reveals germline selection against severe mtDNA mutations. Science 319, 958–962.

Fleischmann, Z., Annis, S., Franco, M., Oreshkov, S., Popadin, K., Woods, D.C., Tilly, J.L., Khrapko, K., 2021. Nuclear DNA-encoded fragments of mitochondrial DNA (mtDNA) confound analysis of selection of mtDNA mutations in human primordial germ cells (preprint). Genomics. 10.1101/2021.10.18.464832

Fleischmann, Z., Cote-L’Heureux, A., Franco, M., Oreshkov, S., Annis, S., Khrapko, M., Aidlen, D., Popadin, K., Woods, D.C., Tilly, J.L., Khrapko, K., 2024. Reanalysis of mtDNA mutations of human primordial germ cells (PGCs) reveals NUMT contamination and suggests that selection in PGCs may be positive. Mitochondrion 74, 101817. 10.1016/j.mito.2023.10.005

Floros, V.I., Pyle, A., Dietmann, S., Wei, W., Tang, W.C.W., Irie, N., Payne, B., Capalbo, A., Noli, L., Coxhead, J., Hudson, G., Crosier, M., Strahl, H., Khalaf, Y., Saitou, M., Ilic, D., Surani, M.A., Chinnery, P.F., 2018. Segregation of mitochondrial DNA heteroplasmy through a developmental genetic bottleneck in human embryos. Nature Cell Biology 2018 20:2 20, 144–151.

Franco, M., Cote-L’Heureux, A., Fleischmann, Z., Chen, Z., Khrapko, M., Vyshedskiy, B., Braverman, M., Popadin, K., Pickett, S., Woods, D.C., Tilly, J.L., Turnbull, D., Khrapko, K., 2023. A novel, wave-shaped profile of germline selection of pathogenic mtDNA mutations is discovered by bypassing a classical statistical bias. 10.1101/2023.11.21.568140

Franco, M., Pickett, S., Fleischmann, Z., Khrapko, M., Annis, S., woods dori Markuzon, N., Turnbull, D., Khrapko, K., 2020. Can detrimental mtDNA mutations be under positive selection in the germline? The FASEB Journal 34, 1–1. 10.1096/fasebj.2020.34.s1.09461

Freyer, C., Cree, L.M., Mourier, A., Stewart, J.B., Koolmeister, C., Milenkovic, D., Wai, T., Floros, V.I., Hagstrom, E., Chatzidaki, E.E., Wiesner, R.J., Samuels, D.C., Larsson, N.-G., Chinnery, P.F., 2012. Variation in germline mtDNA heteroplasmy is determined prenatally but modified during subsequent transmission. Nature Genet 44, 1282–1285. 10.1038/ng.2427

Gunbin, K., Peshkin, L., Popadin, K., Annis, S., Ackermann, R.R., Khrapko, K., 2017. Integration of mtDNA pseudogenes into the nuclear genome coincides with speciation of the human genus. A hypothesis. Mitochondrion 34, 20–23. 10.1016/j.mito.2016.12.001

Gupta, R., Kanai, M., Durham, T.J., Tsuo, K., McCoy, J.G., Kotrys, A.V., Zhou, W., Chinnery, P.F., Karczewski, K.J., Calvo, S.E., Neale, B.M., Mootha, V.K., 2023. Nuclear genetic control of mtDNA copy number and heteroplasmy in humans. Nature 620, 839–848. 10.1038/s41586-023-06426-5

Insalata, F., Hoitzing, H., Aryaman, J., Jones, N.S., 2022. Stochastic survival of the densest and mitochondrial DNA clonal expansion in aging. Proc. Natl. Acad. Sci. U.S.A. 119, e2122073119. 10.1073/pnas.2122073119

Kennedy, S.R., Salk, J.J., Schmitt, M.W., Loeb, L.A., 2013. Ultra-Sensitive Sequencing Reveals an Age-Related Increase in Somatic Mitochondrial Mutations That Are Inconsistent with Oxidative Damage. PLoS Genet 9, e1003794.EP-. doi:10.1371/journal.pgen.1003794

Khrapko, K., Coller, H.A., Andre, P.C., Li, X.C., Hanekamp, J.S., Thilly, W.G., 1997. Mitochondrial mutational spectra in human cells and tissues. Proceedings of the National Academy of Sciences 94, 13798–13803.

Korotkevich, E., Conrad, D.N., Gartner, Z.J., O’Farrell, P.H., 2025. Selfish mutations promote age-associated erosion of mtDNA integrity in mammals. Nat Commun 16, 5435. 10.1038/s41467-025-60477-y

Kraytsberg, Y., Kudryavtseva, E., McKee, A.C., Geula, C., Kowall, N.W., Khrapko, K., 2006. Mitochondrial DNA deletions are abundant and cause functional impairment in aged human substantia nigra neurons. Nature Genet 38, 518–520.

Kremer, L.S., Golder, Z., Barton-Owen, T., Papadea, P., Koolmeister, C., Chinnery, P.F., Larsson, N.-G., 2025. The bottleneck for maternal transmission of mtDNA is linked to purifying selection by autophagy. Sci. Adv. 11, eaea4660. 10.1126/sciadv.aea4660

Kuiper, L.M., Shi, W., Verlouw, J.A.M., Hong, Y.S., Arp, P., Puiu, D., Broer, L., Xie, J., Newcomb, C., Rich, S.S., Taylor, K.D., Rotter, J.I., Bader, J.S., Guallar, E., Van Meurs, J.B.J., Arking, D.E., 2025. Deleterious mitochondrial heteroplasmies exhibit increased longitudinal change in variant allele fraction. iScience 28, 112590. 10.1016/j.isci.2025.112590

Lei, L., Spradling, A.C., 2016. Mouse oocytes differentiate through organelle enrichment from sister cyst germ cells. Science 352, 95–99. 10.1126/science.aad2156

Li, M., Rothwell, R., Vermaat, M., Wachsmuth, M., Schröder, R., Laros, J.F.J., van Oven, M., de Bakker, P.I.W., Bovenberg, J.A., van Duijn, C.M., van Ommen, G.-J.B., Slagboom, P.E., Swertz, M.A., Wijmenga, C., Kayser, M., Boomsma, D.I., Zollner, S., de Knijff, P., Stoneking, M., 2016. Transmission of human mtDNA heteroplasmy in the Genome of the Netherlands families: support for a variable-size bottleneck. Genome Res. 26, 417–426. 10.1101/gr.203216.115

Li, M., Schröder, R., Ni, S., Madea, B., Stoneking, M., 2015. Extensive tissue-related and allele-related mtDNA heteroplasmy suggests positive selection for somatic mutations. also used: Kennedy 84, Radiation Res, 99, 228-48 112, 2491–2496. 10.1073/pnas.1419651112

Liu, Q., Iqbal, M.F., Yaqub, T., Firyal, S., Zhao, Y., Stoneking, M., Li, M., 2022. The transmission of human mitochondrial DNA in four‐generation pedigrees. Human Mutation 43, 1259–1267. 10.1002/humu.24390

Nekhaeva, E., Bodyak, N.D., Kraytsberg, Y., McGrath, S.B., van Orsouw, N.J., Pluzhnikov, A., Wei, J.Y., Vijg, J., Khrapko, K., 2002. Clonally expanded mtDNA point mutations are abundant in individual cells of human tissues. Proceedings of the National Academy of Sciences 99, 5521–5526.

Nicholas, A., Kraytsberg, Y., Guo, X., Khrapko, K., 2009. On the timing and the extent of clonal expansion of mtDNA deletions: evidence from single-molecule PCR. Exp Neurol 218, 316–319.

Otten, A.B.C., Sallevelt, S.C.E.H., Carling, P.J., Dreesen, J.C.F.M., Drüsedau, M., Spierts, S., Paulussen, A.D.C., de Die-Smulders, C.E.M., Herbert, M., Chinnery, P.F., Samuels, D.C., Lindsey, P., Smeets, H.J.M., 2018. Mutation-specific effects in germline transmission of pathogenic mtDNA variants. Hum. Reprod. 33, 1331–1341.

Popadin, K., Polishchuk, L.V., Mamirova, L., Knorre, D., Gunbin, K., 2007. Accumulation of slightly deleterious mutations in mitochondrial protein-coding genes of large versus small mammals. Proceedings of the National Academy of Sciences 104, 13390–13395.

Stewart, J.B., Freyer, C., Elson, J.L., Wredenberg, A., Cansu, Z., Trifunovic, A., Larsson, N.G., 2008. Strong purifying selection in transmission of mammalian mitochondrial DNA. PLoS Biol 6, e10.

Tan, B.G., Mutti, C.D., Shi, Y., Xie, X., Zhu, X., Silva-Pinheiro, P., Menger, K.E., Díaz-Maldonado, H., Wei, W., Nicholls, T.J., Chinnery, P.F., Minczuk, M., Falkenberg, M., Gustafsson, C.M., 2022. The human mitochondrial genome contains a second light strand promoter. Molecular Cell 82, 3646–3660.e9. 10.1016/j.molcel.2022.08.011

Tan, B.G., Wellesley, F.C., Savery, N.J., Szczelkun, M.D., 2016. Length heterogeneity at conserved sequence block 2 in human mitochondrial DNA acts as a rheostat for RNA polymerase POLRMT activity. Nucleic Acids Res 44, 7817–7829. 10.1093/nar/gkw648

Wanrooij, P.H., Uhler, J.P., Simonsson, T., Falkenberg, M., Gustafsson, C.M., 2010. G-quadruplex structures in RNA stimulate mitochondrial transcription termination and primer formation. Proc. Natl. Acad. Sci. U.S.A. 107, 16072–16077. 10.1073/pnas.1006026107

Wei, W., Tuna, S., Keogh, M.J., Smith, K.R., Aitman, T.J., Beales, P.L., Bennett, D.L., Gale, D.P., Bitner-Glindzicz, M.A.K., Black, G.C., Brennan, P., Elliott, P., Flinter, F.A., Floto, R.A., Houlden, H., Irving, M., Koziell, A., Maher, E.R., Markus, H.S., Morrell, N.W., Newman, W.G., Roberts, I., Sayer, J.A., Smith, K.G.C., Taylor, J.C., Watkins, H., Webster, A.R., Wilkie, A.O.M., Williamson, C., Ashford, S., Penkett, C.J., Stirrups, K.E., Rendon, A., Ouwehand, W.H., Bradley, J.R., Raymond, F.L., Caulfield, M., Turro, E., Chinnery, P.F., 2019. Germline selection shapes human mitochondrial DNA diversity. Science 364, eaau6520.

Yuan, Y., Ju, Y.S., Kim, Y., Li, J., Wang, Y., Yoon, C.J., Yang, Y., Martincorena, I., Creighton, C.J., Weinstein, J.N., Xu, Y., Han, L., Kim, H.-L., Nakagawa, H., Park, K., Campbell, P.J., Liang, H., Consortium, P., 2020. Comprehensive molecular characterization of mitochondrial genomes in human cancers. Nature Genetics 52, 342–352.

Zhang, H., Esposito, M., Pezet, M.G., Aryaman, J., Wei, W., Klimm, F., Calabrese, C., Burr, S.P., Macabelli, C.H., Viscomi, C., Saitou, M., Chiaratti, M.R., Stewart, J.B., Jones, N., Chinnery, P.F., 2021. Mitochondrial DNA heteroplasmy is modulated during oocyte development propagating mutation transmission. Sci. Adv. 7, eabi5657. 10.1126/sciadv.abi5657

